# Distinct gene regulatory networks govern hematopoietic and leukemia stem cells

**DOI:** 10.1101/2024.09.28.614980

**Authors:** Boyang Zhang, Alex Murison, Angelica Varesi, Naoya Takayama, Liqing Jin, Nathan Mbong, Erwin M. Schoof, Stephanie Xie, Amanda Mitchell, Julia Etchin, A. Thomas Look, Mathieu Lupien, Mark D. Minden, Jean C.Y. Wang, Peter W. Zandstra, Elvin Wagenblast, John E. Dick, Stanley W.K. Ng

## Abstract

The underlying gene regulatory networks (GRN) that govern leukemia stem cells (LSC) in acute myeloid leukemia (AML) and hematopoietic stem cells (HSC) are not well understood. Here, we identified GRNs by integrating gene expression (GE) and chromatin accessibility data derived from functionally defined cell populations enriched for HSC and LSC. We analyzed n=32 LSC+ and n=32 LSC-cell fractions from n=22 AML patients, along with n=7 stem and n=10 progenitor enriched cell populations sorted from human umbilical cord blood (hUCB), producing a database of n≈17,000 transcription factor (TF) regulatory interactions for hUCB-HSPC and AML. We developed an iterative algorithm that associates the degree of chromatin openness with TF binding preferences, and the GE of candidate TF and target genes within 100kb upstream of transcription start sites. A putative regulatory structure was found to be enriched in HSC-enriched cell populations, comprising TF-target gene interactions between ETS1, EGR1, RUNX2, and ZNF683 oriented in a self-reinforcing configuration. A regulatory loop comprising FOXK1 and MEIS1, rather than the 4-factor HSC subnetwork, was detected in the LSC-specific GRN. The core HSC and LSC TF networks were extended using protein-protein interaction (PPI) data to determine connectivity with interacting genes whose expression strongly associated with LSC/HSC frequency estimates, producing a database of n=103,516 PPI target pathways. The effect of perturbing genes along the identified pathways on functional HSC and LSC frequency was predicted based on statistical regression analyses. To validate GRN predictions, we used pharmacologic and CRISPR targeting, in addition to re-examining published functional data associated with several network nodes that were predicted to impact stemness. Notably, we found that inhibition of CDK6 in AML samples markedly reduced LSC numbers as assessed in de novo serial xenotransplantation studies (fold change ≈ 10), as predicted by the LSC GRN model. Additionally, in-house CRISPR-based knockdown of ETS1 resulted in a significant decrease in HSC quiescence-associated microRNA-126 expression, and increased HSC frequency. Taken together, our models provide a comprehensive view of the underlying regulatory structures governing functional human HSC and LSC. This approach has translational potential as it can be used as a high-throughput in-silico screening tool for the systematic identification of gene targets for LSC elimination and HSC expansion.

## Introduction

A major challenge in the search for functional HSC and LSC regulators can be attributed to the variability in stem cell surface proteome, which makes it difficult to isolate these rare cells at high purity for analysis. The gold-standard assay for the identification of functional stem cells is serial xenotransplantation, which has proven to be a useful tool for probing stem cell biology and identifying molecular factors that govern stemness properties. In fact, several studies involving serial xenografts have demonstrated that perturbing key molecular control points governing stemness can impact functional HSC and LSC frequency^1–4^. Although several key stemness regulators have already been identified, there likely exists many more factors that have yet to be found.

One way to identify additional stem cell regulators is by gaining a better understanding of the regulatory circuitry underlying stemness. In one study, it was found that in mobilized human peripheral blood (hPB) and bone marrow (hBM), CD34^+^ hematopoietic stem and progenitor cells (HSPC) from healthy donors were governed by a core GRN. Specifically, it was found that a heptad of TFs (i.e., SCL, LYL1, LMO2, GATA2, RUNX1, ERG, FLI-1) frequently occupied numerous genomic loci in these cells^5,6^. In CD34^+^ cells isolated from primary AML patient samples, these same factors have been found to promote GE patterns that resembled non-leukemic CD34^+^ HSPC, and was also significantly associated with patient outcome, as well as leukemic engraftment of LSC+ AML patient samples in NOD/SCID mice^7^. A major limitation of using multi-ChIP-Seq profiling as done in these studies to define GRNs is that only a select number of factors were studied. Thus, it is likely that many other components of the GRNs governing HSC and LSC function have yet to be elucidated.

Although there have been clear successes in the identification of stem cell targets, a high-throughput genome-wide method for elucidating GRNs that govern rare functional stem cells in primary human samples is missing. In particular, there is currently no systematic way to rapidly identify additional molecular targets to perturb stemness. To address this, we developed a GRN construction algorithm that integrates GE, chromatin accessibility, and PPI data with functional HSC/LSC frequency estimates derived from primary human samples. We extracted stem cell specific GRN structures, and a comprehensive genome-wide catalogue of candidate HSC/LSC targets. We validated the model predicted effects of perturbing several gene targets (i.e., CDK6 and ETS1) on functional LSC and HSC in-vivo and in-vitro, respectively. The in-silico network models derived provide a principled approach for rapidly screening for gene targets that are likely to impact functional stem cells, thereby accelerating the discovery of novel strategies for targeted LSC eradication and HSC expansion.

## Materials and methods

### Patient samples

All AML samples were collected with informed consent according to procedures approved by the Research Ethics Board of the University Health Network (UHN; REB# 01-0573-C) and viably frozen in the PM Leukemia Bank. LSC+ and LSC− status was scored using the methods described elsewhere^8^. hUCB cells were obtained from consenting healthy donors according to the procedures approved by the Research Ethics Board of the University Health Network and HSC were processed as described previously^2^. CD34^+^ adult hBM was purchased from Lonza (cat. no. 2M-101C). Only hBM obtained from donors between 25 and 35 years of age were used in this study. CD34^+^ hFL cells were purchased from AllCells (AllCells LLC, Alameda, CA, US, cat. no. FL-CD34-001F or FL-CD34-002F/∼20-22wk gestation).

### Identification of functional stem cell molecular features

Isolation of functional human HSCs able to generate a multilineage graft at single-cell resolution has not yet been achieved, with the highest purity being ∼1 in 10 Lin-CD34+ CD38-CD45RA-CD90+ CD49f+ (HSC-enriched) hUCB cells. Consequently, the GE generated from even the most purified samples likely reflects programs of non-HSCs^9^. Similarly, in AML, the surface proteome of blast cells including LSCs has been reported to be highly variable/aberrant between and within patients. Additionally, functional LSC frequency ranges from 1 in a few hundred to millions of cells depending on the patient sample and mouse model used. Consequently, even the GE generated from samples highly enriched for functional LSCs likely reflects programs of non-LSCs^10^. However, HSC- and LSC-specific GE should theoretically correlate with HSC and LSC frequencies as estimated by limiting dilution analysis (LDA) in xenograft experiments, respectively, providing a way to extract features associated with functional stem cells within heterogeneous cell populations.

### Gene expression data processing and analysis

The procedure for RNA-sequencing (RNA-Seq) data generation for hFL and hBM has been previously described^11^. HSC and progenitor subsets from hFL and hBM were sorted (2000-4000 cells) and RNA was isolated using the PicoPure RNA extraction kit. RNA was subjected to amplification using SMARTerTM Ultra Low RNA Kit for Illumina Sequencing (Clontech, Cat. No. 634826). Library preparation of cDNA was performed using the Nextera protocol (Illumina, Nextera DNA Sample Preparation Kit, Cat No. FC-121-1031). Four samples per lane of sequencing on the Illumina HiSeq 2000 platform was configured.

Processing of microarray and RNA-Seq data derived from normal HSPC and AML samples was performed. Briefly, raw Illumina HumanHT-12 V3.0 expression beadchip GE data derived from n=9 hUCB-HSC enriched cell populations (i.e., n=3 LIN^-^CD34^+^CD38^-^CD45RA^-^THY1^+^CD49F^+^, n=3 LIN^-^CD34^+^CD38^-^CD45RA^-^THY1^-^CD49F^+^, n=3 LIN^-^CD34^+^CD38^-^CD45RA^-^THY1^-^CD49F^-^) were downloaded from the Gene Expression Omnibus (GEO) at accession code GSE29105. The fluorescence intensity profiles were subjected to variance stabilization and quantile normalization using the lumi 2.26.4 R package^12^. All data was put into the log base-2 scale. Limiting dilution estimates of the frequency of HSC with long term repopulating potential associated with each GE profile was also obtained^9^. Reads per kilobase of transcript per million mapped reads (RPKM) normalized RNA-Seq data derived from n=12 hFL and n=12 hBM HSC-enriched cell populations (i.e., for each n=3 LIN^-^CD34^+^CD38^-^CD45RA^-^THY1^+^CD49F^+^, n=3 LIN^-^CD34^+^CD38^-^CD45RA^-^THY1^-^CD49F^+^, n=3 LIN^-^CD34^+^CD38^-^CD45RA^+^THY1^-^, n=3 LIN^-^ CD34^+^CD38^-^CD45RA^-^THY1^-^CD49F^-^) along with the corresponding estimates of HSC frequency at limiting dose were obtained from the authors of the associated study ^11^. A value of 1 was added to the RPKM values before applying a log-transformation to the base-2 scale. LSC frequency estimates and GE data derived from LSC+ and LSC-cell fractions were obtained and processed as described previously ^8^. For all GE datasets, the feature with the maximum mean GE for each gene in the data set was used for downstream analyses.

### ATAC-seq data processing and analysis

BWA 0.7.2 software was used to align single-end sequencing reads to the hg19 reference genome, producing ATAC-Seq BAM files corresponding to n=17 HSPC and n=64 AML samples ^13^. MACS 2.1.1 peak calling software (with signal per million reads correction) was used to call significantly enriched peaks and compute fold enrichment of signal versus local background^14^. Further data processing was done based on other work^15–17^. Briefly, BAM files were sorted based on genomic coordinates using samtools 0.1.19. Duplicate reads were filtered using picard 2.9.0-1. Furthermore, reads that were unmapped, mapped to poorly alignable regions^18^, mapped to chromosome Y, or mapped to mitochondrial DNA were removed using samtools. Additionally, only reads with high alignment/mapping quality scores were retained (i.e., Q30) using samtools. Blacklisted regions as defined in the ENCODE mapability consensus excludable BED file and a custom created file ^16^ known to exhibit a high number of artefactual reads in CHIP and ATAC-seq assays were removed. Specific to the current study, a BED file containing the transcription start sites (TSS) of all RefSeq genes from the hg19 assembly was downloaded from the UCSC genome browser^19^. A list of non-overlapping 200bp regions centered on n= 342,857 unique peak summits within +/-100kb flanking the TSS of n=26,966 genes across n=17 HSPC and n=64 AML ATAC-Seq profiles were identified and merged into a single data matrix. Overlapping 200bp windows were merged and new non-overlapping 200bp bins were computed. This above described procedure was performed to filter for regions with high regulatory potential and to reduce the dimensionality of the genome-wide dataset. Raw counts of DNA fragments for each sample that overlapped the n= 342,857 unique 200bp regions was then calculated using bedops 2.4.26. The EDAseq 2.8.0 R package was then used to normalize the data between and within samples^20^. Differential chromatin accessibility between stem and non-stem profiles was evaluated using the voom function included in the limma 3.30.13 R package^21,22^. The normalized data matrix was subsequently partitioned into sub-matrices per gene for downstream analyses. Bedtools 2.19.1 was used to extract FASTA sequences from each open chromatin region for motif analysis to identify potential TF binding sites (TFBS). The motif occurrence detection suite (MOODS) 1.9.3 command-line software package^23,24^ was used with the CISBP 1.02 database of n=868 human TF PWMs^25^ to calculate log-odds binding scores for each open chromatin region in each ATAC-seq profile.

### Core gene regulatory network inference

Regularized linear regression as implemented in the glmnet 2.0-13 R package was used with an L1 penalty while enabling leave-one-out cross validation (LOOCV) to identify a minimal subset of open chromatin regions that may regulate a given target gene of interest within 100kb ^26^. The regions are selected such that the normalized read counts can be combined in a weighted linear fashion to produce overall scores that exhibit coordinated changes with the target GE being modelled. Here, each chromatin region included in the linear model is weighted by a regression coefficient, where the regions associated with the highest positive and negative coefficients are predicted to be the strongest activators and repressors of the target gene of interest, respectively.

To visualize the core TF networks as directed graphs, the qgraph 1.4.3 and igraph 1.0.1 R packages were used. Tarjan’s algorithm^27^ as implemented in the RBGL 1.50.0 package was used to detect strongly connected components (i.e., subsets of factors that potentially regulate each other) within the overall TF-TF regulatory networks. Centrality measures of outdegree and betweenness were computed using the igraph package to identify highly interactive TFs within the network based on their connectedness with other factors. Here, the default settings were used to compute the shortest paths between TFs, where the absolute value of all edge weights were used as input to Dijkstra’s algorithm^28^.

Regularized linear regression using a ridge penalty ^26^ as implemented in the glmnet R package was used while enabling LOOCV to model the relationship between the core TFs and estimates of stem cell frequency. Overall scores were computed using a linear combination of the GE of core nodes weighted by the estimated regression coefficients. The presence of monotonic relationships between the scores and stem cell frequency was subsequently assessed using Spearman correlation analysis, where a statistically significant trend was detected based on a P value of less than 0.05. Core TF networks with scores that are significantly associated with stem cell frequency are predicted to contribute to the regulation of stem cells.

### Core network extension using protein interaction data

Protein-protein interactions (PPI) as defined in the Human Integrated Protein-Protein Interaction rEference (HIPPIE) 2.0 database^29,30^ was used to interrogate the potential connectivity between non-TF genes of interest and the core HSC and LSC TF networks. An edge weight is assigned to each link between node pairs in the extended networks. These values are used as distance metrics during graph traversal to calculate the shortest paths between start and end nodes. Specifically, the values are encoded as the inverse of the PPI quality scores from the HIPPIE database multiplied by the mean GE of node pairs connected by each edge. The inverse operation is used such that network edges, representing PPI between the most abundant gene products, with the strongest experimental evidence is assigned the lowest cost to traverse. Since the edge weights assigned are always positive, the shortest path defined by interacting protein pairs in the HIPPIE database between each query gene of interest and each core TF was evaluated using Dijkstra’s algorithm^28^ as implemented in the igraph R package. The visNetwork 2.0.1 R package was used to visualize the extended networks.

Regularized linear regression using a ridge penalty^26^ as implemented in the glmnet R package was used while enabling LOOCV to model the relationship between the shortest PPI paths from the query genes to the core TFs and estimates of stem cell frequency. Overall pathway scores were computed using a linear combination of pathway nodes weighted by the estimated regression coefficients. The presence of monotonic relationships between the scores and stem cell frequency was subsequently assessed using Spearman correlation analysis, where a statistically significant trend was detected based on a P value of less than 0.05. PPI paths with scores that are significantly associated with stem cell frequency are predicted to contribute to the regulation of stem cells. Components of such PPI paths that have positive or negative coefficients indicates that the associated genes in the model contributes positively or negatively to the overall pathway score, which would be predicted to increase or decrease stem cell frequency, respectively. Thus, perturbation of key PPI path nodes, whose expression is also significantly associated with stem cell frequency, are predicted to change the overall score and impact stem cell supportive signaling, maintenance, function, and/or survival. Control points can be evaluated in both the HSC and LSC models to predict toxicity to healthy and leukemic cells, respectively.

### Computational pipeline for HSC and LSC GRN construction

A computational workflow for GRN model construction was developed that integrates chromatin accessibility (ATAC-Seq, ^16^) and GE data (RNA-Seq and/or microarray) derived from primary hUCB-HSPC and AML patient samples. Specifically, n=17 GE ^31^ and ATAC-Seq profiles for HSC, MPP, MLP, CMP, GMP, MEP, as well as n=32 LSC+ and n=32 LSC-GE and ATAC-Seq profiles^32^ were analyzed together to produce a directed graph. Each genome-wide ATAC-Seq profile contained between 10,000 and 100,000 open chromatin regions. A database of n=868 TFs from the Catalog of Inferred Sequence Binding Preferences (CIS-BP) v1.02 was used to seed network construction^25^. To filter for accessible chromatin regions that have high potential for regulating target genes, the open chromatin regions within 100kb flanking the transcription start site (TSS) of each seed gene was considered for analysis. This window size was chosen since chromatin conformation studies have shown that regions within this range have among the highest probability of making contact with target promoter regions through chromatin looping^33^. This procedure resulted in between 10 to 150 potential regulatory regions per gene. To further narrow down the list of regulatory regions, only differentially open chromatin peaks between at least one pair of cell types was considered for further analysis (i.e., LSC+, LSC-, HSC, MPP, MLP, common myeloid progenitor (CMP), GMP, monocytes, and granulocytes). Motif analysis using the CIS-BP v1.02 database was performed on the exposed nucleotide sequences corresponding to the resulting subset of open chromatin regions to predict TF binding at each locus, identifying potential regulatory relationships between the TFs in the full CIS-BP database. As shown by others, higher motif scores frequently indicate more likely TF binding events, as confirmed using ChIP-Seq assays^34,35^. Sparse linear regression was used to select a minimal number of open regions around each gene’s TSS whose degree of openness across cell-type specific ATAC-Seq profiles can explain the fluctuations in GE of each target gene^26^. The regions assigned the highest positive and negative standardized regression coefficients were predicted to indicate loci of binding events with the strongest activating and inhibitory effects on target GE, respectively. This approach limits the high number of potential regulatory links for each target gene and allows for the construction of parsimonious GRNs. TFs corresponding to the top 10 motifs that are predicted to bind to these regions are then scored based on: 1) overall level of target GE, 2) degree of openness of region, 3) whether the direction of potential regulator and target GE corresponds to the sign (i.e., positive or negative) of the regression coefficient, 4) whether the potential regulator is expressed when the region is open, 5) whether the potential regulator’s GE is enriched in stem or non-stem cells, 6) whether the potential regulatory region is more accessible in stem or non-stem cells, and 7) correlation of regulator GE with estimates of stem cell frequency. Based on these criteria, the most likely regulatory factors are predicted to bind to the selected open regions and regulate target GE, as reflected by the edge thickness in the resulting directed graph. TF-TF regulatory interactions that are enriched in stem and non-stem cells are stored for further analysis. This process is iterated to infer additional regulatory relationships among the TFs in the CIS-BP database until a strongly connected component (i.e., SCC, a directed graph where each node is reachable by all other nodes) is identified or when no new regulatory links are found (Figure 1). Identification of SCCs in the network may indicate an important set of TFs that regulate one another in a given cellular compartment. Depending on which genes are in the SCCs, and which cell-type their regulatory interactions are enriched in, SCC members may represent important nodes for maintaining a particular cell-type specific GE state. Such nodes may be recognized as vertices with high betweenness or outdegree measures of centrality, due to forming densely interconnected sub-networks comprising other genes with known cell-type specific functions. SCCs therefore might serve as critical regulatory hubs that if repressed or reinforced, may disconnect or strengthen many regulatory paths between other nodes in the network, respectively, which would likely lead to changes to GE state that may affect cellular function. By using the GRN construction workflow to interrogate all open chromatin regions within the flanking 100kb regions of all gene TSS’s in the transcriptome for which data was available (n≈8,000), we generated databases of n≈17,000 TF-target gene regulatory interactions specific to human HSPC and AML, while marking interactions that were predicted to be HSC/LSC-specific. These exploratory databases include a list of target genes that are potentially regulated by the n=868 human TFs with DNA binding preference data available in the CIS-BP v1.02 database. Taken together, our GRN construction algorithm can produce predictions of directed gene-gene interactions, which in turn can be used to build directed graphs similar to other studies that successfully used DNase-Seq data^36^ to recapitulate well studied GRNs in muscle cells and embryonic stem cells (ESC) ^37^.

**Figure 1:**
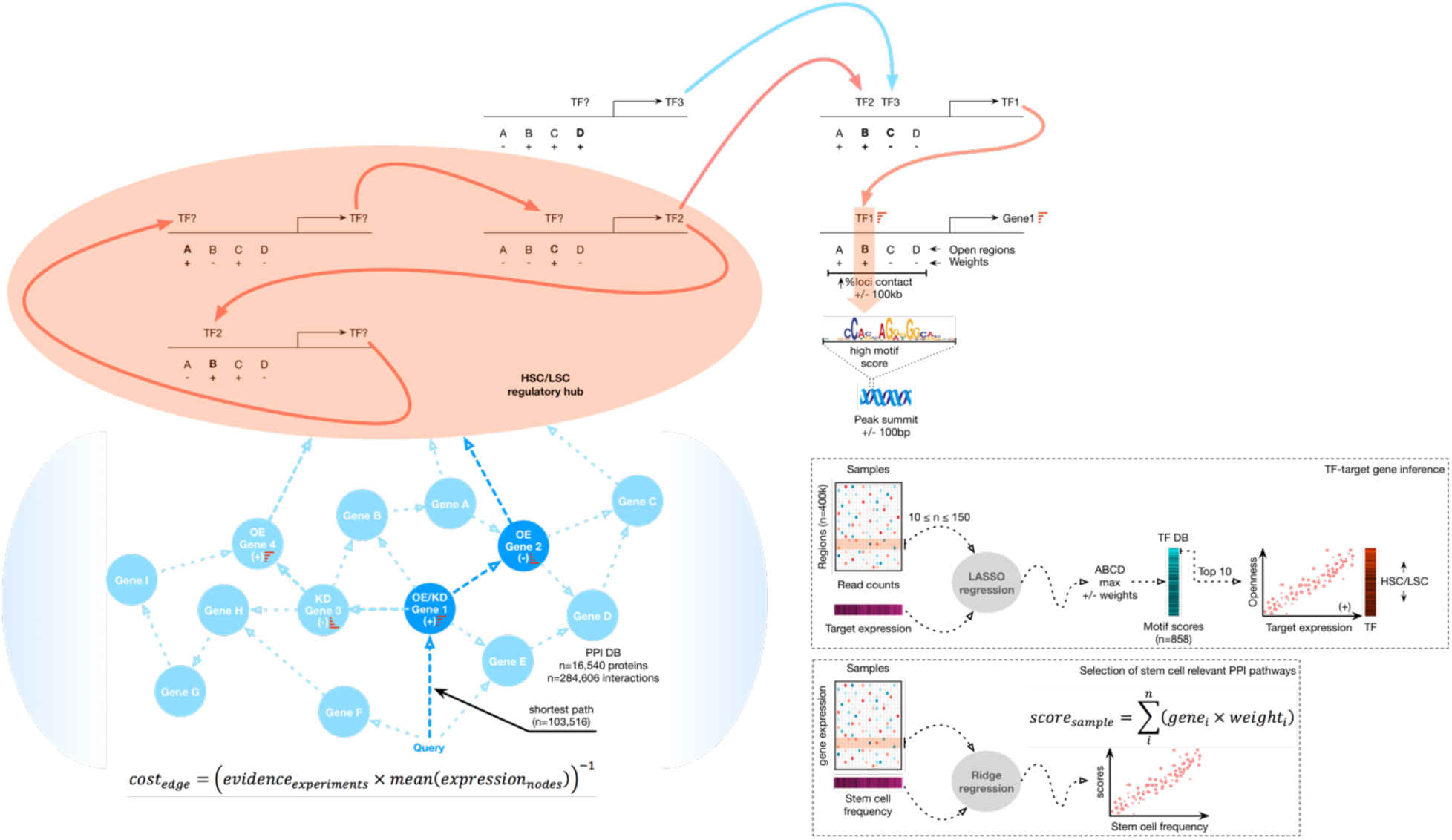
Overview of GRN inference and target gene identification. Top: Conceptual schematic illustrating the inference of core self-sustaining TF networks. Hypothetical accessible chromatin regions A, B, C, and D that may regulate target genes upon TF binding. Plus (+) and minus (-) signs corresponding to red and blue arrows together indicate activation and repression of target GE due to TF binding, respectively. The pink highlighted subset of TFs maintains each other by forming a positive feedback loop. Bottom left: Hypothetical example depicting the process of using protein interaction data to extend core HSC/LSC GRNs through the identification of interacting gene products. Boxes: Regression-based workflows illustrating the integration of GE and ATAC-Seq data with TF binding scores and HSC/LSC frequency to infer TF-TF physical interactions (top), and the selection of HSC/LSC-relevant PPI pathways by associating its GE scores to stem cell frequency (bottom).

### Selection of GRN-predicted KD and OE stemness targets

Targets that can induce a decrease in LSC while increasing HSC upon KD or OE would inform novel therapeutic strategies in AML. Specifically, the OE candidates are genes whose expression is positively and negatively correlated to HSC and LSC frequency, while also having positive and negative weights in the HSC and LSC PPI pathway linear regression models, respectively. In this way, OE of such candidates in the HSC and LSC models would have the effect of increasing and decreasing the overall pathway scores, respectively, since a positive and negative component of the score is amplified in the respective models. Conversely, the KD candidates are genes that have correlation patterns and regression coefficient signs that are opposite to the potential OE targets. Thus, KD of these candidates would also lead to increasing and decreasing the overall pathways scores in HSC and LSC models, respectively, since a negative and positive component of the score is nullified in the respective models. Lastly, the PPI pathways with which the potential OE and KD targets are associated have GE scores that are positively correlated to both HSC and LSC frequency. Therefore, perturbations that increase and decrease HSC and LSC scores would be predicted to increase and decrease HSC and LSC frequency, respectively. Of note, CDK6 is ranked as the second most important KD target in this category based on the number of PPI paths that it is associated with.

Targets that can induce a decrease in LSC while having minimal effects on HSC upon KD or OE are also important candidates for the development of novel AML treatments. In this case, the OE candidates are genes with expression that is negatively correlated with LSC frequency and have negative weights in the LSC PPI pathway models that have GE scores that are positively correlated to LSC frequency. Accordingly, OE of these candidates would have the effect of lowering the overall pathway scores since a negative component of the score is amplified. Ultimately, a lower pathway score would be predicted to be associated with lower LSC frequency. To be considered a potential OE target in this category, the expression of the gene and the associated HSC PPI pathway GE scores must not be correlated to HSC frequency. The KD candidates, on the other hand, follow the above criteria, except that they are associated with GE that is positively correlated with LSC frequency, and have LSC PPI pathway model weights that are positive. Thus, KD of these candidates would also lead to a lower overall pathway score since a positive component of the score is nullified, which would be predicted to lower LSC frequency. In this category of functional stem cell KD targets, XPO1 was ranked second.

Finally, there is translational interest to expand HSC to allow broader use in HSC transplant therapies. Thus, perturbing targets that can induce an increase in HSC even though LSC are minimally affected upon KD or OE is worthwhile, as the former can potentially gain a competitive advantage over the latter. In this category of potential pro-HSC targets, OE candidates are defined to have GE and associated HSC PPI pathway model scores that are positively correlated with HSC frequency, while having positive weights in the models. At the same time, potential OE targets must not have GE or LSC PPI pathway model scores that are significantly correlated with LSC frequency. In this way, OE of these candidates in the HSC model would lead to an increase in the overall pathway scores since a positive component of the score is amplified, which would be predicted to promote the HSC state. The KD candidates follow the same criteria as the OE targets but have GE that is negatively correlated with HSC frequency, and have negative weights in the associated HSC PPI pathway models. Thus, KD of these candidates would also lead to a higher overall pathway score since a negative component of the score is nullified, which would be predicted to reinforce the HSC state.

### Analysis of LSC scores using the NanoString assay

We submitted the 104 Affymetrix probeset identifiers corresponding to LSC-associated genes, along with reference genes chosen to cover a wide range of expression levels in AML^38^, to NanoString Technologies^39^ for custom codeset creation. The 100-base pair (bp) NanoString probes were fabricated to overlap or be proximal to the corresponding Affymetrix probe target regions. In each lane of the NanoString cartridge used, 150 ng of RNA per sample (5 μl) was incubated with 20 μl of reporter probe and 5 μl of capture probe mix (supplied by the manufacturer) at 65 °C for 16 to 24 h for hybridization on the nCounter Prep Station (version 4.0.11.1). After hybridization, excess probes were washed out using a 2-step magnetic bead-based purification strategy according to the manufacturer’s protocol, and purified target/probe complexes were immobilized on the NanoString cartridge for data collection. Transcript counts were determined using the nCounter Digital Analyzer (version 2.1.2.3) at the high-resolution setting. Specifically, digital images were processed with final barcode counts tabulated in reporter code count (RCC) output files.

The NanoString assay was performed using RNA from bulk mononuclear AML patient cells obtained from A. Thomas Look and Julia Etchin. RCC files containing raw transcript counts from each cartridge were analysed using the nSolver analysis software (version 2.0.72) for quality control (QC) and normalization purposes using default settings for GE analysis. Specifically, RCC files for each cartridge along with a reporter library file containing codeset probe annotations were imported into nSolver. The software was used to normalize the captured transcript counts to the geometric mean of the reference genes included in our assay and the codeset’s internal positive controls, and to check for imaging, binding, positive spike-in, and normalization quality. The output files from nSolver were read into R for further QC, normalization, and data processing. The normalized GE counts were log2-transformed after incrementing by 1. The LSC17 score, along with the Spearman correlation to the average GE of the 104 LSC-associated genes in LSC+ and LSC-cell fractions was computed for each GE profile corresponding to KPT-8602 or vehicle treated AML samples using the scaled data.

### CDK6 inhibitor experimental procedure

Animal experiments were done in accordance to the institutional guidelines approved by the University Health Network animal care committee. NSG mice (NOD.Cg Prkdc^scid^Il2rgtm1Wjl/SzJ; Jackson Laboratory) were sub-lethally irradiated with 250 rads 24 hours before intrafemoral injection with AML patient cells. Xenografts were treated at 4 weeks post-transplantation with vehicle control (PBS with 0.5% methylcellulose) or PD0332991 (100mg/kg dissolved in vehicle) by oral gavage for 14 days and then euthanized for primary transplant analysis. The injected femur and non-injected bones (tibias and contra-lateral femur) were collected and crushed separately in Iscove’s modified Dulbecco’s medium and human AML chimerism was assessed with the following antibodies with flow cytometry on a BD Celesta (from BD, note need catalog numbers): V500-anti-CD45, BV768-anti-CD33, PE-anti-CD117, APC-anti-CD33, FITC-anti-CD15, PECy5-anti-CD14 and sytox blue (ThermoFisher, S34857, 1:3000) for cell viability. Human leukemic cells (CD45+CD33+) used for cell cycle analysis and limiting dilution secondary transplantation studies were isolated from primary grafts by cell sorting on FACS AriaIII that had previously been viably frozen in the following manner: 1-3 mouse xenografts were thawed and pooled to make two technical replicates that were then depleted of mouse cells via a mouse depletion kit (Miltenyi, according to manufacturer’s instructions) on LS columns (miltenyi) using MACS magnet techology and then stained with FITC-anti-CD45, APC-anti-CD33 and propidium iodide. Isolated cells from 3 patient AMLs treated with vehicle or drug were pooled from the technical replicates and injected at defined doses in NSG mice and euthanized at 8 weeks post-transplantation to assess for presence of AML grafts. Injected femur and non-injected bones were isolated and flushed separately and analyzed on a BD Celesta with the same antibody cocktail as for primary grafts. A mouse was considered engrafted if CD45+CD33+ was >0.05%. LSC frequency was estimated using Elda software (http://bioinf.wehi.edu.au/software/elda/; ^40^. AML cells isolated from primary xenografts from four patient samples were fixed and stained for Ki67-Hoechst assays as described ^2^ for cell cycle status on a BD Fortessa.

### Cord blood lineage depletion and sorting

Human cord blood samples were obtained from Trillium and William Osler hospitals with informed consent in accordance to guidelines approved by University Health Network (UHN) Research Ethics Board. Individual cord blood samples were pooled and processed within 24-48 hours after birth. Lineage depleted cells were isolated with the StemSep Human Hematopoietic Progenitor Cell Enrichment Kit (StemCell Technologies, 14056) and Anti-Human CD41 TAC (StemCell Technologies, 14050) according to the manufacturer’s protocol.

To sort LT-HSCs, up to 1x10^7 lineage depleted cells were stained in 1ml with the following antibodies (all from BD Biosciences, unless stated otherwise): CD45 V500 (10ul, 560777, HI30), CD34 APC-Cy7 (5ul, clone 581, custom conjugation), CD38 PE-Cy7 (5ul, 335790, HB7), CD90 APC (20ul, 559869, 5E10), CD45RA FITC (20ul, 555488, HI100) and CD49f PE-Cy5 (20ul, 551129, GoH3). LT-HSCs were sorted on the FACSAria III (BD Biosciences) based on CD45+CD34+CD38-CD45RA-CD90+CD49f+^9^.

### CRISPR knockdown experimental approach

Electroporation of LT-HSCs: Sorted LT-HSCs were cultured for 36-48 hours in serum-free X-VIVO 10 (Lonza) media with 1% Bovine Serum Albumin Fraction V (Roche, 10735086001), 1X L-Glutamine (Thermo Fisher, 25030081), 1X Penicillin-Streptomycin (Thermo Fisher, 15140122) and the following cytokines (all from Miltenyi Biotec): FLT3L (100ng/mL), G-CSF (10ng/mL), SCF (100ng/mL), TPO (15ng/mL) and IL-6 (10ng/mL). Electroporation of chemically synthesized gRNAs and Cas9 protein was carried out. In short, gRNAs were synthesized from IDT as Alt-R CRISPR/Cas9 crRNAs. Both crRNAs against control, ETS1 or EGR1 were combined with tracrRNA in a single tube to anneal at 95^°^C for 5 min. For each reaction, 1.2µl crRNA:tracrRNA, 1.7µl Cas9 protein (IDT) and 2.1µl PBS were combined and incubated for 15 min at room temperature. 1µl of 100µM electroporation enhancer (IDT) was added. Pre-cultured LT-HSCs were resuspended in 20µl of Buffer P3 (Lonza) per reaction and added to the ribonucleoprotein complex. LT-HSCs were electroporated with the Lonza Nucleofector (V4XP3032) using the DZ-100 program. Cells were recovered overnight at 37^°^C by adding 180 µl of X-VIVO media.

gRNA design: gRNAs were designed with the CRoatan algorithm^41^. ETS1 gRNAs target exon 8 and result in a drop-out of 200 bp. EGR1 gRNAs target exon 2 and result in a drop-out of 300 bp. Control gRNAs target exon 1 of OR2W5 and result in a drop-out of 150 bp. gRNA sequences: Control gRNA-1: GACAACCAGGAGGACGCACT, Control gRNA-2: CTCCCGGTGTGGACGTCGCA, ETS1 gRNA-1: TTACCTCCAGGTAAACTCGG, ETS1 gRNA-2: TGTCCTTATTGAGGTCAGCA, EGR1 gRNA-1: ACTCCTTCAGCGCCTCCACA, EGR1 gRNA-2: CCAGGAGCGATGAACGCAAG.

Assessing CRISPR/Cas9 efficiency: Genomic DNA was isolated using Agencourt GenFind V2 (Beckman Coulter, A41499) ^42^. The CRISPR/Cas9 edited locus was amplified by PCR using the AmpliTaq Gold 360 Master Mix (ThermoFisher, 4398881). Subsequently, PCR products were purified using the ZR-96 DNA Clean-up Kit (Zymo Research, D4018) and submitted for Sanger sequencing using the reverse PCR primer. The chromatograms were analyzed using the online tool ICE (https://ice.synthego.com) in order to assess CRISPR/Cas9 efficiency. PCR primers: Control (OR2W5) forward primer: 5’-TCGGCCTGGACTGGAGAAAA-3’, Control (OR2W5) reverse primer: 5’-GAGACCACTGTGAGGTGAGA-3’, ETS1 forward primer: 5’-TTCCTCAGAGTAAACCCATGACC-3’, ETS1 reverse primer: 5’-GAGTGGCTGGAGAGAGACTAAAG-3’, EGR1 forward primer: 5’-CATGATCCCCGACTACCTGT-3’, EGR1 reverse primer: 5’-CATCTGACCTAAGAGGAACCCT-3’.

### qRT-PCR for miR-126

After electroporation of LT-HSCs with control, ETS1 or EGR1 gRNAs, cells were cultured in serum-free X-VIVO 10 (same as above) for 5 days. miR-126 expression profiling was carried out using the TaqMan MicroRNA Cells-to-CT Kit (Thermo Fisher, 4391848). Quantitative PCR analysis was performed on the Roche Lightcycler 480. All signals were quantified using the ΔCt method and were normalized to the levels of RNU48. qPCR probes: hsa-miR-126: Assay ID 002228, RNU48: Assay ID 001006.

### Animal studies

The University Health Network (UHN) Animal Care Committee approved all mouse experiments. All xenotransplantations were performed with 8- to 12-week-old female NOD.Cg-PrkdcscidIl2rgtm1Wjl/SzJ (NSG) mice (JAX) and male NOD.Cg-PrkdcscidIl2rgtm1WjlTg(CMV-IL3,CSF2,KITLG)1Eav/MloySzJ (NSGS). Both mouse strains were sublethally irradiated with 225cGy 24 hours prior to transplantation. Sample size was chosen to give sufficient power for calling significance with standard statistical tests. 150 CRISPR/Cas9 edited LT-HSCs were intrafemorally injected as previously described^43^. After 20 weeks, primary transplanted mice were sacrificed to obtain the right femur (RF) and bone marrow (left femur and both tibias, BM). Bones were flushed with PBS + 2.5% FBS and counted using the Vicell XR (Beckman Coulter). A small subset of cells was frozen down for genomic DNA isolation. Only mice that showed a CRISPR/Cas9 knock-out efficiency of >90% as determined by Sanger sequencing and a CD45+ engraftment level in the RF of >5% were utilized in the analysis. After 9 weeks, secondary transplanted mice were sacrificed to obtain the right femur (RF) and left femur (LF). Again, bones were flushed, counted using the Vicell XR and a subset of cells were frozen down for genomic DNA.

Flow cytometric analysis: Cells from transplanted mice were stained with the following antibodies (all from BD Biosciences, unless stated otherwise): CD45 APC-Cy7 (1:100, 348795560566), CD45 A700 (1:100, 560566), CD33 APC (1:100, 340474) and CD19 V450 (1:100, 560353), CD41 PE-Cy5 (1:200, Beckman Coulter, 6607116), GlyA PE (1:100, Beckman Coulter, IM2211U) and CD3 FITC (1:100, 349201). Cells were analyzed on the FACSCelesta (BD Biosciences).

Secondary transplantations: For secondary transplantations, RF and BM samples were thawed and pooled. Subsequently, murine cells were depleted using the mouse cell depletion kit (Miltenyi Biotec, 130104694). After depletion, cells were stained as described above for cord blood sorting. Human CD45+ cells were sorted using the FACSAria III (BD Biosciences) and injected at different doses into irradiated NSGS mice. Stem cell frequency was estimated using the online tool ELDA (http://bioinf.wehi.edu.au/software/elda/index.html) ^40^. The threshold for detection of human engraftment was 0.1% CD45+ cells in the RF.

### Protein mass spectrometry

Data generated by ultra-high-performance liquid chromatography (UHPLC) coupled with high resolution mass spectrometry was analyzed to investigate the protein expression levels of CDK6 and cyclin D3 in n=26 LSC+/- cell fractions sorted from primary patient samples. Details regarding the mass spectrometry workflow are detailed elsewhere in full^44^.

### Statistical analysis

The R 3.3.2 statistical programming environment was used for all statistical analyses. The Spearman rank-order correlation coefficient is evaluated to determine the presence and strength of monotonic relationships between covariates unless specified otherwise. A P value threshold of 0.05 is used to determine statistical significance unless specified otherwise.

## Results

### Development of a GRN construction algorithm that accounts for xenograft estimates of functional stem cell frequency

In order to systematically identify TFs that are likely to be essential to functional HSC and LSC, we set out to define GRN models by integrating ATAC-Seq and GE data on highly purified populations of human HSCs and progenitors, as well as AML populations sorted into LSC+ and LSC-fractions. In all cases, we used xenograft transplantation assays to estimate the frequency of HSC and LSC in each sorted fraction (Figure 1, Supplemental information).

### Elucidating a gene regulatory network governing human HSC

The GRN construction algorithm (see methods) was applied to the hUCB-HSPC ATAC-seq/GE, and functional HSC frequency data. An analysis of the resultant HSC network was performed to detect the presence of SCC that took the form of positive feedforward regulatory loops that may be important for the maintenance of HSC specific programs. A core network of n=6 TFs was identified (i.e., FOSL1, ZNF274, EGR1, ZNF683, RUNX2, and ETS1), which formed two interconnected positive feedforward regulatory loops, where GE and chromatin peaks were highly enriched in the HSC compartment (Figure 2a). The 4-factor sub-circuit comprising EGR1, RUNX2, ZNF683, and ETS1, in particular, was found to have relatively higher betweenness centrality measure than other factors in the network, with ETS1 having the highest value (Figure 2b). These genes act as intermediate GRN nodes in the shortest path between many other TFs, suggesting that they may be regulated by many factors. In endothelial cells, ETS1 has been found to directly regulate miR-126, which has been reported to govern the balance between self-renewal and quiescence in human HSC^3^. Specifically, high miR-126 levels were associated with increased HSC quiescence. Indeed, CRISPR-mediated knockdown of ETS1 resulted in a ≈2-fold decrease in miR-126 (p=0.0002), ≈3.6-fold increase in HSC frequency (p=0.002), and increased CD34+/CD38-/CD90+ cells, as well as human CD45^+^ BM engraftment in xenografts (Figures 4e-f and S1). Furthermore, several predicted ETS1 target TFs have been associated with HSC maintenance and function according to various studies (NR4A3^45^, LCOR^46^, FOSB^47^, FOXO4^48^, FLI1^49^, HOXB3/4^50^, EGR1^51^, IRF1^52^). In support of this, similar results were observed in extended experiments with EGR1 (Figures 4e-f). Thus, our results suggest that ETS1 is an important regulator of human HSC.

**Figure 2:**
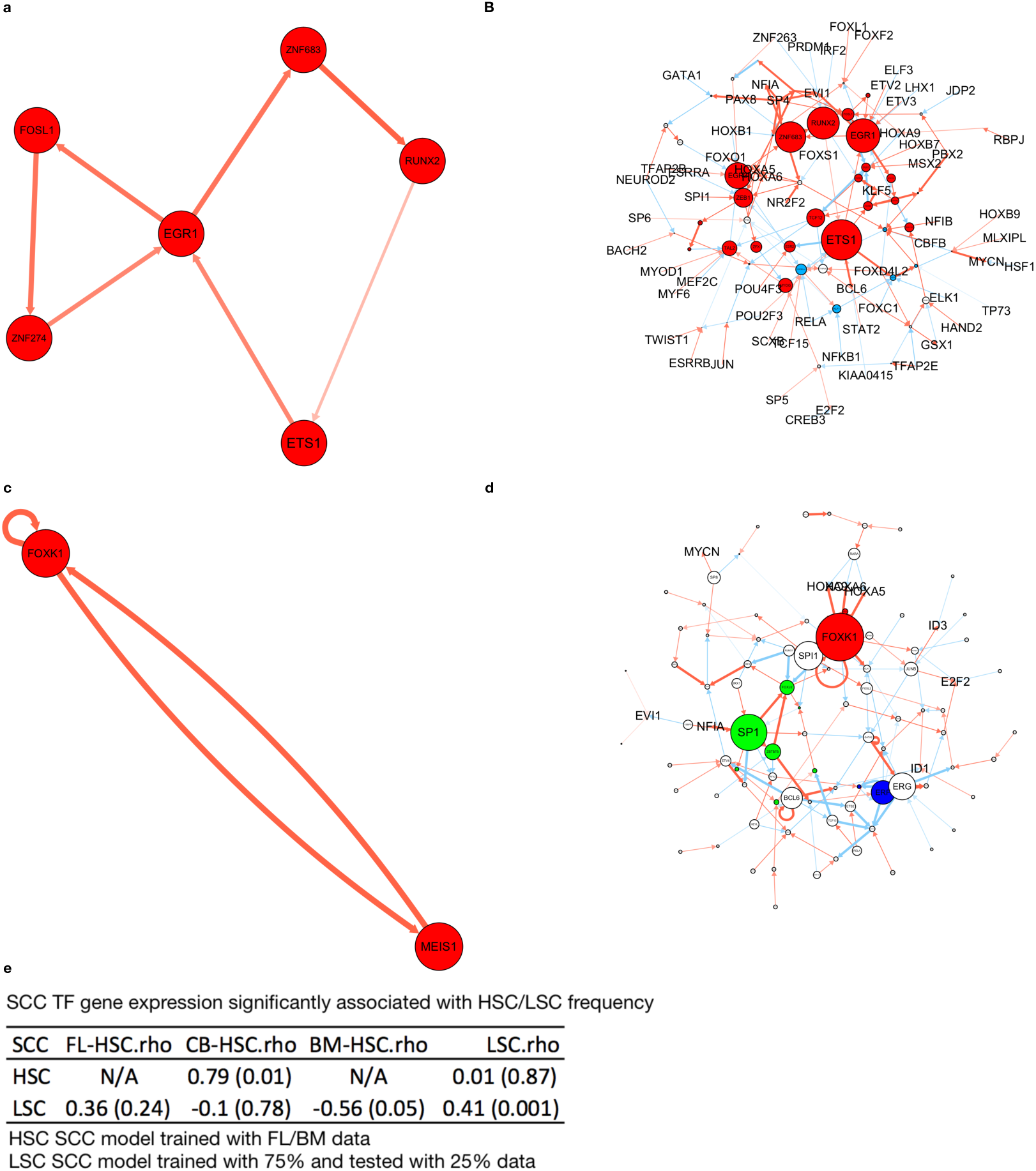
GRNs associated with HSC and LSC. **a,** Two interconnected positive feedforward loops of 6 nodes forms a core HSC TF network. **b,** The core HSC TF network nodes within the overall GRN have high betweenness centrality measures. **c,** A positive feedforward loop of MEIS1 and FOXK1 forms a core LSC TF network. **d,** The core LSC TF network nodes within the overall GRN. FOXK1 has the highest outdegree centrality measure. Nodes are colored according to SCC membership in each respective cell-type specific GRN. In all panels, red and blue curves denote activation and inhibition of target gene transcription, respectively. **e,** Spearman correlation with P-Values in brackets between TF GE and HSC/LSC frequency estimates.

We next reasoned that if the core TFs identified were relevant to functional HSC, then it should be possible to derive a linear model based on their GE to explain hFL/hBM-HSC frequency data. Indeed, training such a model using regression using data from previous work ^11^, produced GE scores that were significantly correlated with HSC frequency estimates of independent samples ^9^ (Figure 2e). These results provide support for the relevance of the GRN-identified TFs to functional human HSCs, demonstrating the feasibility of our approach. In sum, our results suggest that inhibiting core HSC GRN TFs may disrupt functional HSC transcriptional programs. Notably, no positively-reinforced feedforward loops were found in the progenitor compartment, perhaps owing to grouping of MLP, CMP, GMP, and MEP profiles together for collective analysis due to the limited sample numbers of each respective progenitor cell type.

### Elucidating a gene regulatory network governing human LSC

The GRN derivation workflow was next applied to data derived from 32 LSC+ and 32 LSC-sorted cell fractions from AML patients, resulting in an overall AML regulatory network. An SCC in the form of a positive feedforward regulatory loop between MEIS1 and FOXK1 was detected, where both the GE and chromatin peaks were highly enriched in LSC+ samples (Figure 2c). Of note, no such regulatory structures were found in the LSC-compartment. MEIS1 has been reported to promote the LSC state in MLL-driven AMLs^53^, but it is not known whether this TF is also important to LSC in other AML subtypes. Nevertheless, MEIS1 transcript and protein levels are both higher in LSC+ compared to LSC-fractions across a range of high-risk AML samples of diverse cytogenetic backgrounds (P < 0.001 and rank-sum P = 0.01, respectively).

FOXK1 was found to have the highest outdegree centrality measure in the overall network (Figure 2d), suggesting that this TF may be binding to open chromatin around many other TFs, potentially regulating them. FOXK1 has been reported to drive proliferation, invasion, and metastasis in tumors of the ovary^54^, colon^55^, brain^56^, and stomach ^57^, enabling tumor cells to seed malignant growth in new areas of the host. FOXK1 was also reported to interact with FHL2, the loss of which leads to decreased HSC self-renewal and quiescence under conditions of replicative stress such as that induced by serial transplantation^58^. Given that FOXK1 is significantly higher expressed in LSC+ compared to LSC-fractions (P = 0.01), it is possible that this factor may play a role in LSCs as well. Moreover, several predicted FOXK1 target TFs have been associated with cancer or stem cell maintenance and function (HOXA4/5/13^59–62^, RUNX1^63,64^, HMGA1^65^, FOXD1^66^, FOXC2^67^, TGIF2^68^, MEF2C^69^).

The GE of the core LSC network nodes in 75% of the LSC+/LSC-AML cell fractions data was used to derive a linear model to explain the differences in LSC frequencies estimated across different samples in the remaining 25% of the dataset. The resulting GE scores were found to be significantly correlated with LSC frequency, providing support for the relevance of the GRN-identified TFs to functional LSCs (Figure 2e).

### Extending the core HSC and LSC networks using protein interaction data enables rapid in-silico screening for stem cell relevant gene targets

To investigate how additional query genes of interest could be connected to the TFs in the HSC and LSC core networks, Dijkstra’s algorithm^28^ was applied to the HIPPIE protein interaction database to identify the shortest PPI paths from each query gene to each core TF ^29,30^ (Figure 1). Linear regression was then used to relate the GE of nodes along each PPI path to HSC and LSC frequency estimates. PPI paths with GE scores that were found to be significantly associated with HSC and/or LSC frequency were considered for further analysis. This network extension strategy was applied using an initial list of n=83 candidate genes for which an association with HSC or LSC frequency has been observed to explore their connectivity to the core TF networks (Table 2). This query gene list includes receptors associated with enhanced self-renewal in HSC-enriched cell populations^70–72^, miR-126 targets within the PI3K/AKT/MTOR pathway that was shown to play a role in HSC/LSC quiescence and self-renewal^3,4^, LSC genes^8^, and genes that are commonly mutated in clonal hematopoiesis and preleukemic HSCs^73–76^.

### GRN models accurately predicts the effects of perturbing HSC and LSC regulators

To determine the relevance of the identified HSC/LSC relevant genes within the GRNs constructed, we tested whether perturbing key nodes would affect HSC and LSC function. We started with CDK6 as we previously reported a role for this gene in regulating the timing of quiescence exit in hUCB-HSCs^2^ (Figure 3a). Whereas CDK6 OE in HSCs induces faster cell-cycle entry from the G0 state and therefore more self-renewal divisions per unit time, CDK6 inhibition with the PD033299 (PD) compound induces prolonged quiescence. In agreement with this, CDK6 GE is negatively correlated with hUCB and hBM HSC frequency. This makes sense since HSC have low CDK6 GE and protein levels, allowing them to remain quiescent (Figure 3b). Furthermore, CDK6 has a negative regression coefficient in the HSC model (Figure 3a).

**Figure 3:**
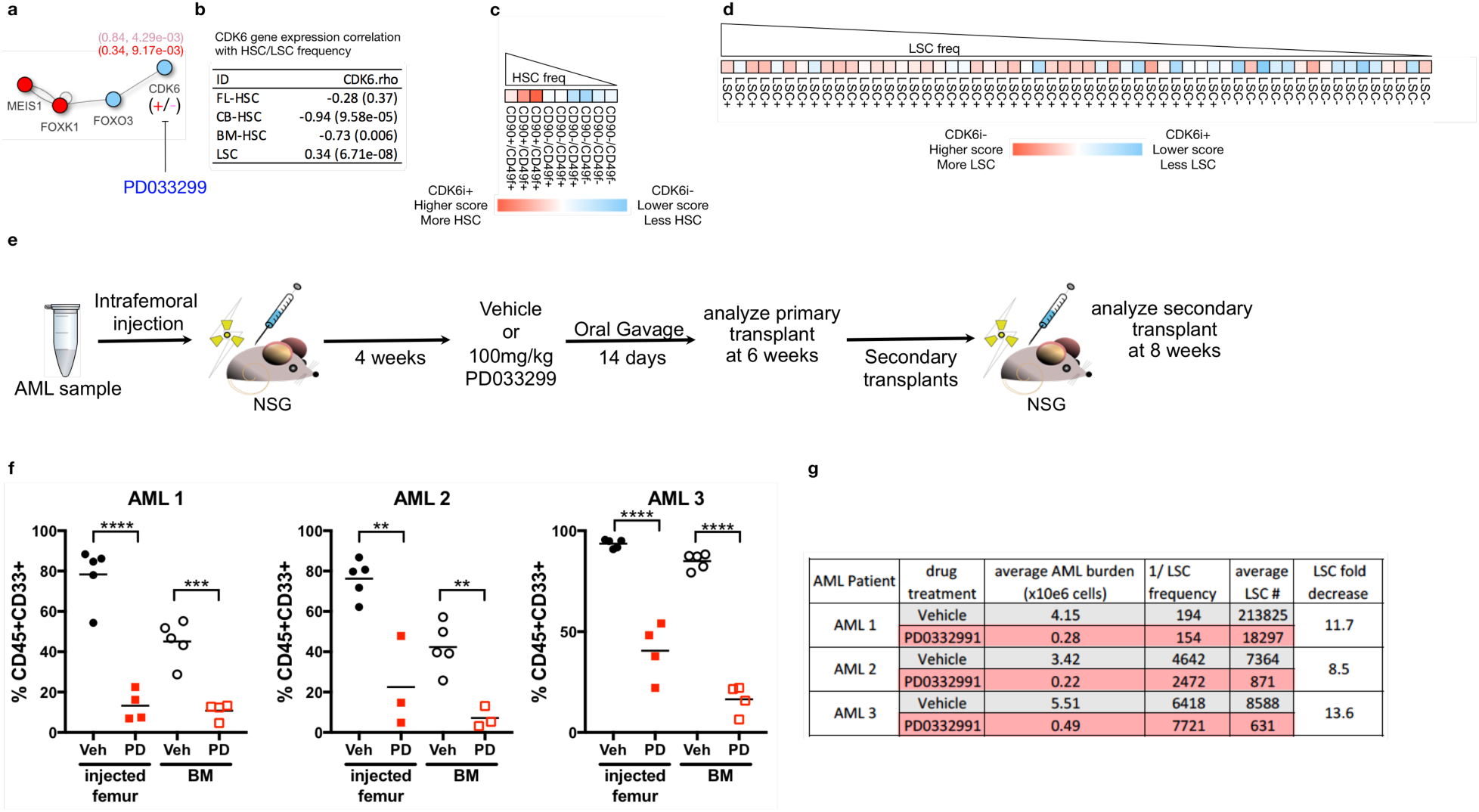
CDK6 inhibition is predicted to reduce LSC with minimal toxicity to HSC. **a,** PPI path containing CDK6 is inhibited by small molecule PD033299, reducing a positive and negative component of the overall pathway activity score in LSC and HSC, respectively. **b,** CDK6 GE correlation with HSC and LSC frequencies. **c,d,** Heatmaps showing the relationship between PPI pathway activity scores and HSC and LSC frequencies. **e,** Schematic of the approach for serial xenotransplantation experiments. **f,** Primary AML engraftment in immune-deficient mice is significantly reduced upon inhibiting CDK6 compared to controls. **g,** Change in LSC frequency after normalizing to the number of cells surviving after treatments.

Thus, decreasing its GE has the effect of raising the overall PPI pathway activity scores in the HSC model, which would be predicted to correspond to an increase in the number of quiescent HSCs (Figure 3c), which is consistent with results from CDK6 inhibitor studies^2^. Conversely, CDK6 expression is positively correlated with LSC frequency (Figure 3b), indicating that LSCs may be more primed for earlier cell cycle entry, which may provide them with an ability to outcompete their normal more quiescent HSC counterparts. CDK6 also has a positive regression coefficient in the LSC GRN (Figure 3a). Therefore, decreasing its GE lowers the overall PPI pathway activity scores, which would be predicted to result in decreased LSC numbers (Figure 3d). To validate this prediction, we carried out serial xenotransplantation studies (Figure 3e). We found that PD-mediated CDK6 inhibition in several primary AML patient samples induced a marked reduction in primary leukemic engraftment (i.e., ≈15-fold reduction) (Figure 3f). In serially transplanted recipients, we found that the overall LSC frequency in the PD and control conditions to not be significantly different (P > 0.05), suggesting that PD treatment eliminated bulk blasts as well as functional LSC to a similar degree, as no LSC enrichment was observed (Figure 3g). This outcome is different from treatment with the standard 7+3 agents that are commonly given to AML patients, which typically lead to higher LSC frequency upon relapse^77^. After accounting for the drastic reduction in leukemic blasts after PD treatment compared to controls, a ∼10-fold decrease in the estimated number of LSC became evident in secondary recipients (Figure 3g). A potential mechanism for PD-mediated LSC elimination is related to decreases in antioxidants that was reported to be caused by CDK6 inhibition in CDK6 and cyclin D3 expressing cells. In such cells, high levels of reactive oxygen species can induce apoptosis in several tumor types^78^. Accordingly, we next assessed CDK6 and cyclin D3 protein levels using protein mass spectrometry for n=26 LSC+/- cell fractions sorted from primary patient samples.

We found cyclin D3 protein levels to be significantly associated with LSC frequency (r = 0.48, P = 0.01). Indeed, cell fractions with the highest LSC frequencies also had the highest levels of cyclin D3 protein (within 1.7 standard deviations (sd) from the maximum level of the n≈8000 proteins detected). Furthermore, we also found that CDK6 protein levels were similarly high in LSC+ cell fractions (within 1.5 sd from the max protein levels in our dataset). As the AML samples interrogated using PD inhibitors are considered to have high LSC frequencies (Figure 3g), they likely express high levels of CDK6 and cyclin D3. Thus, it is possible for PD treatment to induce LSC cell death. Notably, cells remaining after PD treatment were observed to be highly proliferative, suggesting that these residual cells may be sensitized to anti-proliferative standard chemotherapeutic agents.

We previously reported that miR-126 targets many components within the PI3K/AKT/MTOR pathway in human hUCB-HSCs to regulate the balance between quiescence and self-renewal^3^. To investigate how miR-126 targets may be associated with the GRN identified in HSC data, the PPI analytic workflow was applied. In the resulting extended HSC network, it was found that 3 out of 3 (100%) miR-126 targets that connected to the core HSC GRN TFs had positive regression coefficients (Figure 4a). Thus, it follows that miR-126 KD will lead to higher expression of miR-126 target genes, resulting in higher overall PPI pathway activity scores in the HSC model, predicting that there will be more HSC supportive signaling (Figure 4b).

**Figure 4:**
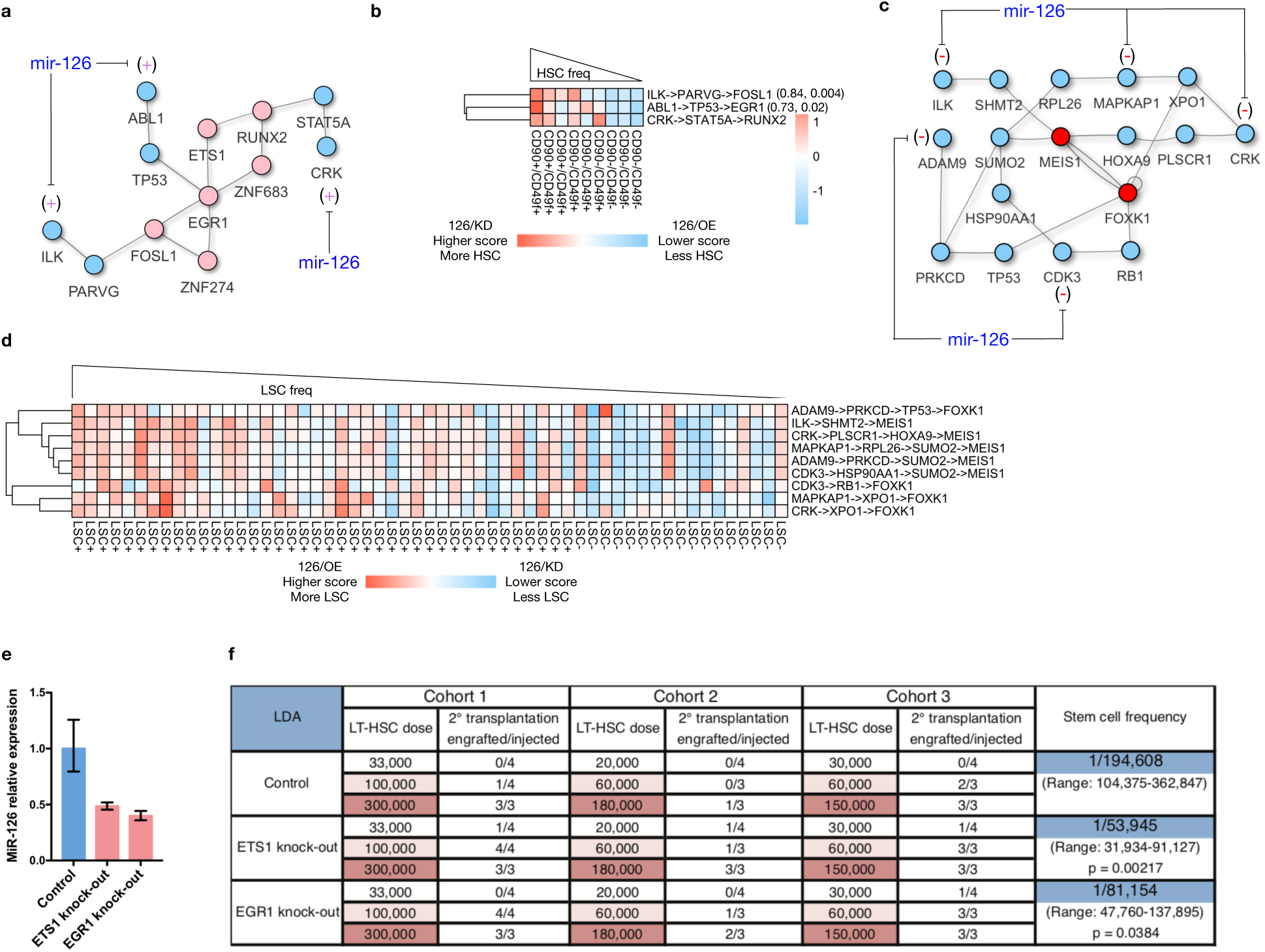
miR-126 knockdown is predicted to reduce LSC and increase HSC. **a,c,** PPI paths containing elements of the PI3K/AKT/MTOR pathway are strongly associated with LSC and HSC frequency. **b,d,** Heatmaps showing the relationship between PPI pathway activity scores and HSC and LSC frequencies. **e,** miR-126 expression as measured by qPCR, and **f,** limiting dose estimates of HSC frequency from secondarily transplanted xenografts, in ETS1 and EGR1 KD versus control conditions in hUCB-HSC.

Conversely, miR-126 OE is predicted by this model to lower HSC numbers. These predictions are consistent with the experimental consequences of perturbing miR-126 in hUCB-HSC, where miR-126 KD and OE results in higher and lower HSC frequency, respectively^3^. On the other hand, in the LSC network, it was found that 9 out of 11 (81.8%) miR-126 targets had negative regression coefficients in the LSC model (Figure 4c). Thus, miR-126 KD and OE would be predicted to lead to higher and lower miR-126 target GE levels, which would contribute to decreasing and increasing the associated PPI pathway activity scores in the LSC model, respectively. Such an effect would be further predicted to result in dampened and heightened LSC activity under the respective conditions (Figure 4d). Indeed, these GRN model predictions reflect the experimental outcomes of perturbing miR-126 in primary AML patient samples and the 8227 AML cell-line. Specifically, miR-126 KD leads to increased proliferation and differentiation of LSC, whereas miR-126 OE leads to enhanced quiescence while favoring self-renewing divisions upon cell cycle transit. Moreover, high miR-126 expression has been observed in relapse and treatment-resistant LSC-enriched primary patient samples^4^.

PROCR (a.k.a., EPCR or CD201) (Figure 5a) is upregulated at the GE and protein levels upon stimulation with the HSC self-renewal inducing UM171 compound, and is a marker of HSCs^70,71^. In our data, PROCR GE was found to be positively correlated with HSC frequency in both hUCB and hBM, suggesting that PROCR may mark HSCs in these tissues (Figure 5b).

**Figure 5:**
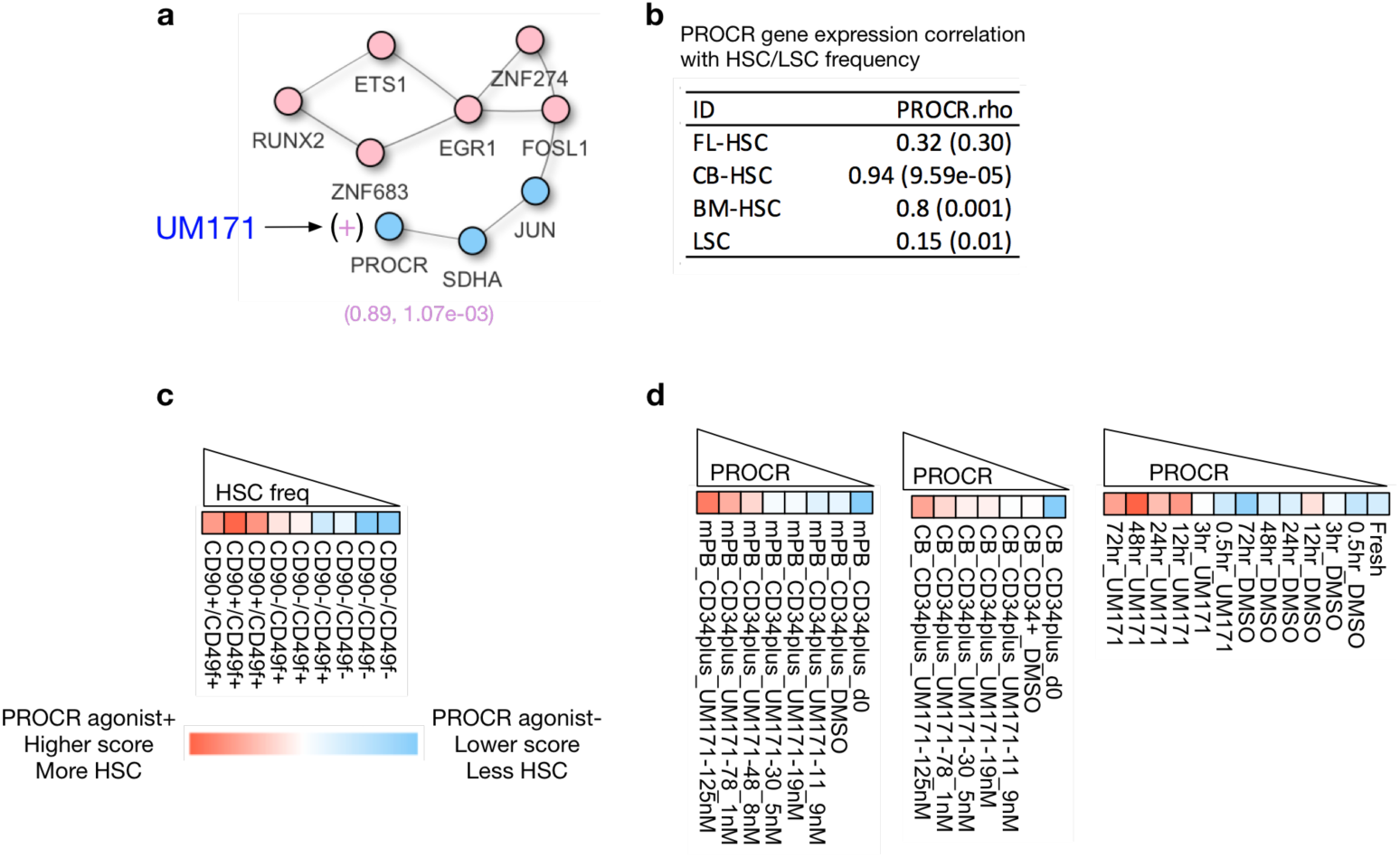
Increased PROCR expression is predicted to increase HSC. **a,** PPI paths containing PROCR are activated by the UM171 compound, increasing a positive component of the overall pathway activity score. **b,** PROCR GE correlation with HSC and LSC frequencies. **c,** Heatmap showing the relationship between PPI pathway activity scores and HSC frequency. **d,** Data from hUCB-HSC expansion studies shows that UM171 treatment induces PROCR GE in a time and dose dependent manner.

Although preliminary work by others identified a subset of PROCR+ hBM cells, it remains unclear whether there was enrichment for HSCs^79^. Since there is a significant positive correlation between PROCR GE and LSC frequency, some LSC may also express this marker (Figure 5b).

We found that PROCR has a positive regression coefficient in the HSC GRN (Figure 5a). Thus, increasing its GE raises the overall activity score of the PROCR PPI pathway, which would be predicted to increase HSC frequency (Figure 5c). These results are consistent with studies that found PROCR KD to result in lower HSC engraftment in vivo ^71^, while self-renewal inducing UM171 treatment increases PROCR GE in CD34+ hUCB/hMPB-HSPC in a time and dose dependent manner (Figure 5d).

Two PPI paths involving XPO1 were identified in the LSC but not the HSC extended GRN, suggesting that perturbation of this node may have LSC-specific effects. XPO1 is responsible for transporting approximately 200 proteins and RNA molecules associated with apoptosis, cell cycle regulation, and tumor suppression from the nucleus to the cytoplasm (Figure 6a). In prior work, treatment with XPO1 inhibitors KPT-330 (Selinexor) and KPT-8602 was found to significantly deplete LSC, supposedly due to a dependency on XPO1-mediated nuclear export, as determined by serial transplantation of primary AML patient samples into immune-deficient mice, while minimal toxicity to normal HSPCs was observed^1,80^. In the current analysis, we found XPO1 GE to be positively correlated with LSC but not HSC frequency (Figure 6b). Since XPO1 has a positive regression coefficient in the extended LSC network, decreasing its GE lowers the overall activity score of the XPO1 PPI paths, which would be predicted to decrease LSC frequency (Figure 6c). These model predictions are consistent with results from the aforementioned XPO1 inhibitor studies^1^. Additionally, two nodes connected to XPO1 are also assigned positive weights in the LSC extended GRN model and have GE that is positively correlated to LSC frequency (i.e., BIVM (r = 0.45, P = 3.87e-13) and PHGDH (r = 0.20, P = 0.002), Figure 6a). In a separate experiment, we used a NanoString assay that implements a set of 104 LSC-associated genes to assess the previously described LSC17 score, which has robust prognostic power in adult and pediatric AMLs^8,81^. In the current study, we also used the data from this assay to calculate scores based on the correlation to the average GE of the 104 genes in LSC+ and LSC-cell populations. The assay was applied to an AML sample with normal cytogenetics and the FLT3-ITD mutation before and after XPO1 inhibition, as described previously^1^. Briefly, this sample was injected into NSG mice that were then treated with the second generation XPO1 inhibitor KPT-8602. Upon BM collection and analysis after 96 hours, we observed a decrease in LSC17 score and correlation to LSC+ GE, concomitant with an increase in the correlation to LSC-GE (Figure 6d). These results further support the LSC GRN prediction that XPO1 inhibition leads to decreased LSC, which is likely to improve AML patient outcomes. Finally, we found that XPO1 PPI pathway scores are not significantly associated with HSC frequency in the HSC GRN model, further predicting that XPO1 inhibitors will not negatively impact normal HSCs. Indeed, this prediction is also consistent with experimental observations from previous reports^1^.

**Figure 6:**
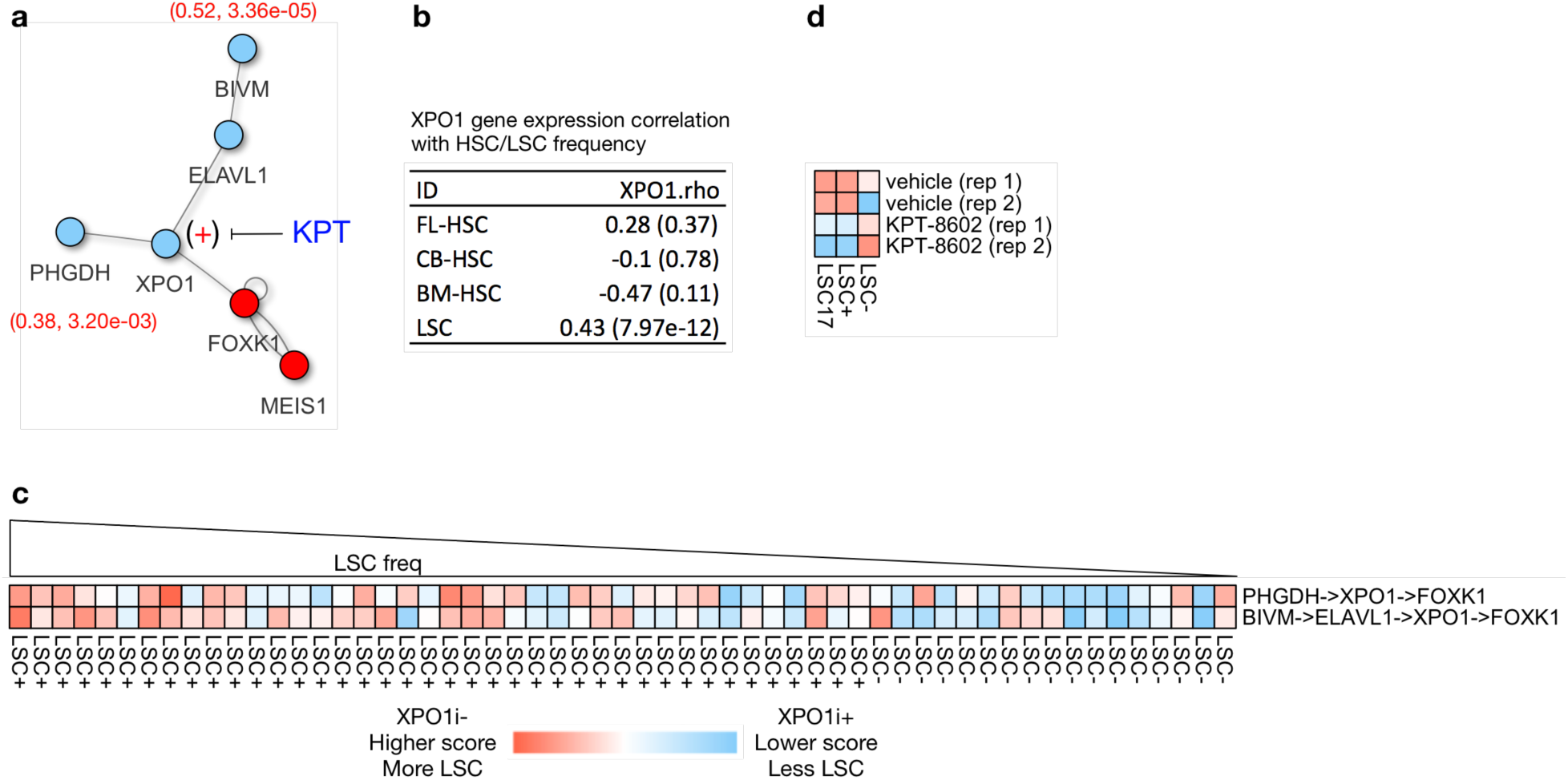
XPO1 inhibition is predicted to reduce LSC with minimal toxicity to HSC. **a,** PPI paths containing XPO1 are inhibited by KPT compounds, reducing a positive component of the overall pathway activity score in LSC. **b,** XPO1 GE correlation with HSC and LSC frequencies. **c,** Heatmap showing the relationship between PPI pathway activity scores and LSC frequency.

### Development of a genome-wide catalogue of gene targets to facilitate rapid systemic identification of functional HSC and LSC modulators

To provide a resource for exploring HSC and LSC relevant GRN targets, we next sought to expand our analysis to determine all potential PPI pathways between all genes in the transcriptome that were captured in our datasets (n≈15,000) and the core HSC and LSC TF GRNs. To accomplish this, we applied our network extension algorithm (see methods) and generated a database of n=103,516 candidate HSC and LSC PPI paths, while predicting the effects of perturbing individual pathway nodes on HSC and LSC frequency. To facilitate target selection, we ranked genes based on the number of PPI pathways with which each had membership, where a higher number of pathways is interpreted as having a higher likelihood of having an impact on functional HSC/LSC. Candidate target genes are assigned to one or more categories based on their predicted effects on HSC and LSC when perturbed, including: 1) OE increases HSC and decreases LSC, 2) KD increases HSC and decreases LSC, 3) OE decreases LSC with negligible impact on HSC, 4) KD decreases LSC with negligible impact on HSC, 5) OE increases HSC with negligible impact on LSC, and 6) KD increases HSC with negligible impact on LSC (Tables S1).

## Discussion

Our study describes a novel computational framework for elucidating GRN structures governing human HSC and LSC. In particular, TF binding site (TFBS) information was integrated with GE, chromatin accessibility, and PPI data to infer regulatory logic. Critically, in-vivo functional readouts of stem cell frequency estimates from xenotransplantation assays were used to guide the extraction of regulatory interactions that were most associated with cell populations enriched with functional HSC and LSC. The compiled databases of TFBS-target gene activation/repression relationships for the HSPC and AML datasets are exploratory resources containing a global map of potential HSC/LSC regulators. We detected minimal subsets of TFs that mutually reinforced the expression of one another in a self-sustaining feed-forward configuration in both HSC and LSC, similar to the regulatory loop found between the pluripotency factors OCT4, SOX2, and NANOG in ESC^82^. Analysis of the PPI-extended HSC and LSC networks revealed that there are likely many pathways that collectively contribute to stemness properties, as opposed to a single pathway being exclusively responsible for maintaining a stem cell state. We therefore assembled a separate database of candidate PPI HSC and LSC targets including their predicted effects on stem cell frequency upon perturbation in each respective cell type, which can also serve as a tool for further probing into the HSC and LSC biology.

Our results demonstrate the feasibility of using GRN models that have been inferred using integrated multi-omics and functional xenograft data to identify gene products that may be essential for HSC expansion or LSC eradication. Our model predictions for CDK6 and ETS1 were validated experimentally using serial xenotransplantation assays, while several other predicted targets agree with previous work^1–4^.

To conclude, we report the first rapid and unbiased in silico screen for probing stemness-relevant pathways and identifying gene targets that are likely to be essential to functional human HSC and LSC. Thus, our results will enhance our understanding of the biology of these stem cells and accelerate the discovery of LSC-directed therapies with minimal toxicity to HSC, as well as inform novel HSC expansion strategies. It is conceivable that stem cell based GRN activity scores could be developed for monitoring treatment efficacy, minimal residual disease status, prognosis, and the emergence of LSC in healthy individuals with clonal hematopoiesis. As it has been reported that stem cell GE programs possess prognostic value in many cancers^83^, it would be of great interest to identify common regulatory structures that are shared across diverse cancer stem cell (CSC) driven tumor types. Such a discovery has the potential to inform the design of pan-CSC-targeted treatments, which would stimulate further application of GRN derivation approaches to investigate the underlying transcriptional circuitry governing diverse pre-malignant and malignant cell populations for vulnerabilities.

## Acknowledgements

We thank Gary Bader, Ruth Isserlin, and Carl White for discussions on GRN inference, network visualization, protein interactions, and PPI databases. We thank Mihai Albu from Tim Hughes lab at the University of Toronto for providing data and scripts for determination of transcription factor binding preferences from the CIS-BP database. We thank The Princess Margaret Genomics Centre (PMGC) for the generation of GE data for the AML samples used in the KPT-8602 inhibitor studies.

## Supplemental Figures

**Figure S1:**
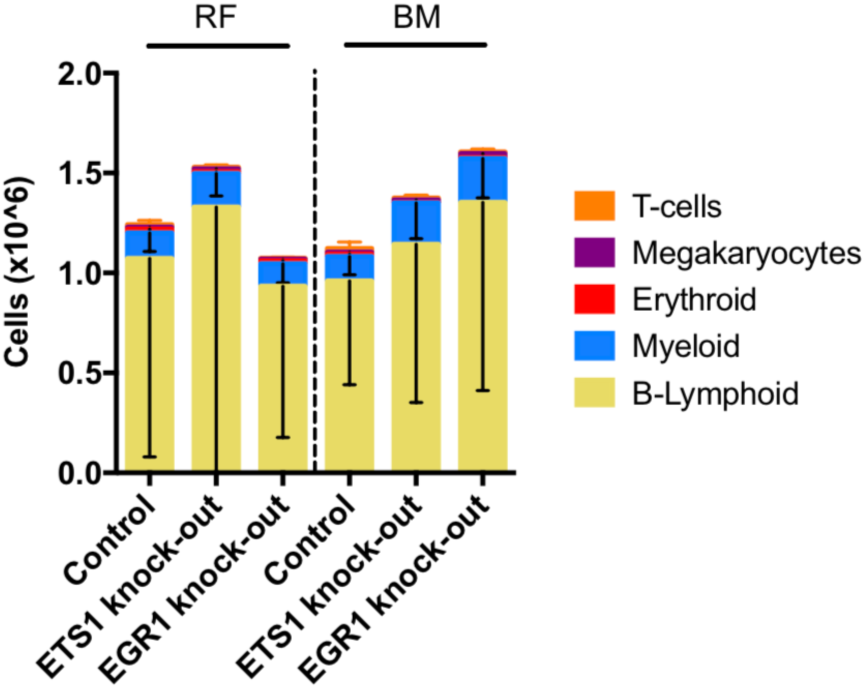
Human CD45+ cell numbers between experimental conditions. Counts of myeloid and lymphoid cells as sorted from secondary xenografts across control and KO conditions.

